# Host use does not drive genetic structure of mountain pine beetles in western North America

**DOI:** 10.1101/2022.06.28.498011

**Authors:** Celia K. Boone, Kirsten M. Thompson, Philippe Henry, Brent W. Murray

## Abstract

The mountain pine beetle (MPB) is one component of an intensively studied co-evolved host-pest system. We investigated the spatial genetic structure of MPB within its historic and recent geographic range expansion as it relates to host use in western North America using 13 pre-selected microsatellite loci. AMOVA shows that genetic structure is not correlated with the host tree species, arguing against the formation of host-race within this species. STRUCTURE analysis shows 4 main clusters in western North America: Northern - Northern British Columbia/Alberta; Central - Southern British Columbia/Alberta/Washington/Idaho/Montana; Southwest - Oregon/California/Nevada and; Southeast - Utah/Wyoming/Arizona/Colorado/South Dakota. Heterozygosity, allelic richness, and number of private alleles is greatest in the Southwest cluster. This cluster correlates with one of the three refugia hypothesized from a recent analysis of neo-Y haplotypes by Dowle and colleagues and represents an important reservoir of MPB genetic diversity.

## Introduction

Ecosystems are exposed to considerable anthropogenic influence, which can lead to regime shifts and changes in species distributions (Vucetich and Waite 2003; Raffa et al. 2008). These adaptations are especially important because they influence species that have lifecycles dependant on environmental inputs (Folke et al. 2004). Insect pests comprise a considerable portion of environmentally sensitive species that are both ecologically and economically important (Kirk et al. 2013). Bark beetles are particularly responsive to changes in climate which can lead to faster life cycles and more generations per year (Werner and Holsten 1985; Bentz and Powell 2014; Six and Bracewell 2015). North America has recently experienced wide-ranging outbreaks of the mountain pine beetle (*Dendroctonus ponderosae* (Hopk.) (Coleoptera: Curculionidae) (MPB), impacting over 26 million ha in Canada alone, with economic impacts expecting to continue for decades as timber supply declines due to the significant quantities of beetle-killed wood (Corbett et al. 2016). While MPB outbreaks are part of a natural cycle facilitating the maintenance of biologically diverse and functionally healthy forest landscapes (Axelson et al. 2009), these substantial impacts, accompanied by range expansion into previously unsuitable habitats, lead to questions about what influences the population structure, and by extension, the genetics, of these insects as part of a broader quest for improved management strategies (British Columbia Ministry of Forests 2013; Fettig et al. 2014).

The mountain pine beetle is a native facultative predatory bark beetle of pine in western North America, with a geographic range extending from the south to Mexico and into the north of British Columbia (BC) and now Alberta (AB) (Mock et al. 2007; Cullingham et al. 2011). MPB outbreaks have been recorded in BC since the early 1900s, following a cyclical pattern with three population phases: endemic, incipient-epidemic, epidemic, and a return to endemic (Wood and Unger 1996; Safranyik and Carroll 2006; Hrinkevich and Lewis 2011). MPB infests several species in the genus *Pinus* in western North America, primarily attacking lodgepole (*P. contorta* var. *latifolia* (Engelm.) Critchfield), ponderosa (*P. ponderosa* Douglas ex C. Lawson), western white (*P. moticola* Douglas ex D. Don), limber pine (*P. flexilis* E. James), and whitebark pines (*P. albicaulis* Engelm.) (Furniss and Schenk 1969; Wood 1982). The most recent major range expansion of MPB began with wind-assisted movement of beetles over the Rocky Mountains in the northwest of Alberta in 2006 (Jackson et al. 2008; Government of Alberta 2014). MPB continues to affect forests in Alberta a decade after the breach of this major geographical barrier and heavy infestations centred in Jasper National Park raise concerns about further expansion (Trevoy et al. 2018). This concern is substantiated by the continuation of MPB’s migration to and establishment at higher elevations in sensitive ecosystems in several states in the western United States (Bentz et al. 2010; Fettig et al. 2014). The cosmopolitan diet and destructive nature of MPB has bolstered interest in the drivers of genetic structure of this beetle in North America.

Influence of host species on the genetics of Dendroctonus has been investigated since the early work on the genus. Initial studies used allozymes to compare the genetic signatures of many species of bark beetle in North America. While variation was detected, many of the studies compared beetles sampled from multiple host tree species from different locations, thus conflating the effect of geography and host species (summarized in Langor and Spence 1991). An allozyme study that sampled declining MPB populations on sites containing two beetle host species found that geographic location was likely a stronger influence than host tree (Langor and Spence 1991). In contrast, Sturgeon and Mitton (1986) compared hosts in geographically proximal sites in Colorado and found an effect of host equal to that of location and contended that host effects may increase genetic heterogeneity and lead to the development of pre-adapted sub-species, or host-races, of MPB. Sampling using amplified fragment length polymorphism (AFLP) markers on MPB from eight sites in the United States and Canada expanded host species comparison and added a landscape genetic structure component, finding a general pattern of isolation-by-distance with little effect of host (Mock et al. 2007). This study, however, combined several host tree species from divergent sites and mixed different trapping methods (below the bark and pheromone-baited traps) which may have obscured the influence of host. Several other studies have assessed landscape genetic structure of MPB using microsatellites (Samarasekera et al. 2012) and single nucleotide polymorphisms (SNPs) (Janes et al. 2014; Batista et al. 2016), finding two major clusters of genetic similarity roughly divided into north and south populations. However host use was excluded from these analyses and only Batista et al. (2016) included sites from the United States.

The effect of MPB host use on genetic heterogeneity leads to two possible hypotheses: one, if host tree species influence MPB genetic structure, we would expect to see more variation between different these species found on the same or similar sites compared to geographic or conspecific groupings; and two, alternatively, if host trees do not influence MPB genetic variation significantly, we would expect to see more variation linked to geographic or conspecific groupings compared to host groupings. Our study uses 13 pre-selected microsatellites to expand upon previous research into the genetic structure of MPB in the United States and Canada. We use a more inclusive dataset of 82 sites in North America to assess general genetic structure, and then create a subset of sites for more detailed comparison of MPB host use during the irruptive phase. We hypothesize that our markers will show a similar north and south clustering compared with that found by Samarasekera et al. (2012), Janes et al. (2014), Batista et al. (2016), but, due to the larger geographic scope of our study, will also show increased sub-clustering in the southern sampling areas.

## Materials and Methods

### Site selection and beetle collection

Beetles used in this study were chosen from a collection procured between 2003 to 2012 from 153 active sites in 105 locations in three provinces in western Canada and eleven states in the western United States (Figure S1). These beetles were primarily collected from lodgepole pine, but also include outbreaks in six other pine species (Table 1). Sample sites for this study were selected based on current and historical mountain pine beetle outbreak activity and include a subset of samples from 2005-2008 analysed by Samarasekera et al. (2012), insects collected by Mock et al. (2007) and previously unreported samples collected from 2010 to 2012 conducted in targeted areas/host tree species not included in previous sampling efforts (Table 1). In prior studies (Samarasekera et al. 2012), western Canada was sampled more intensively than the US. To minimize bias during analysis of genetic structure, 82 sites were selected for a representative geographical distribution throughout western North America. Beetle larvae and adults were collected from individual galleries from multiple trees in each location prior to emergence or from traps during dispersal. A Global Positioning System (GPS) location was determined for each tree sampled or trapping location, and beetles were transported live in bark discs and flash frozen in liquid nitrogen or preserved in 95% ethanol. All samples were stored at -20°C or -80°C prior to DNA extraction. Site locations were visualized using ArcGIS version 10.5.

**Table 1.**
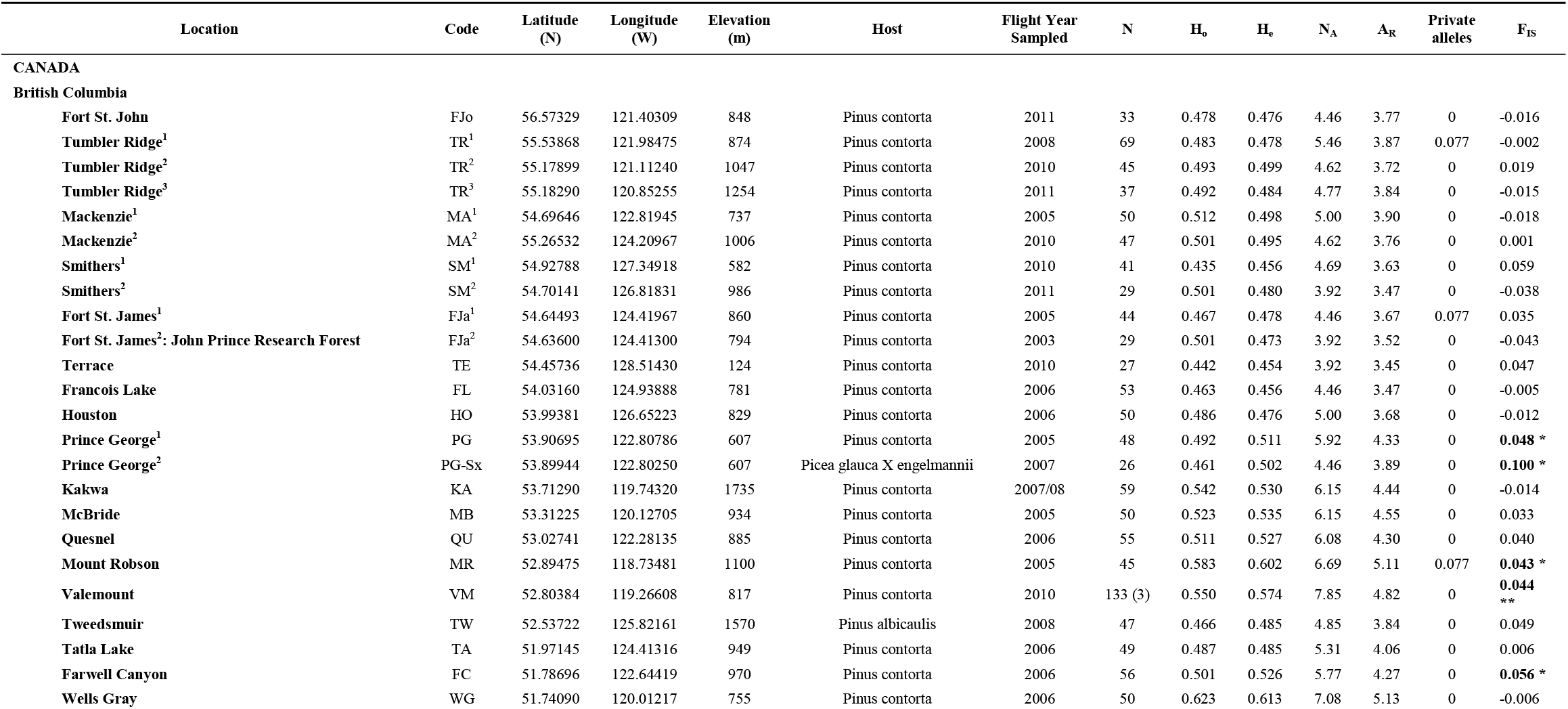
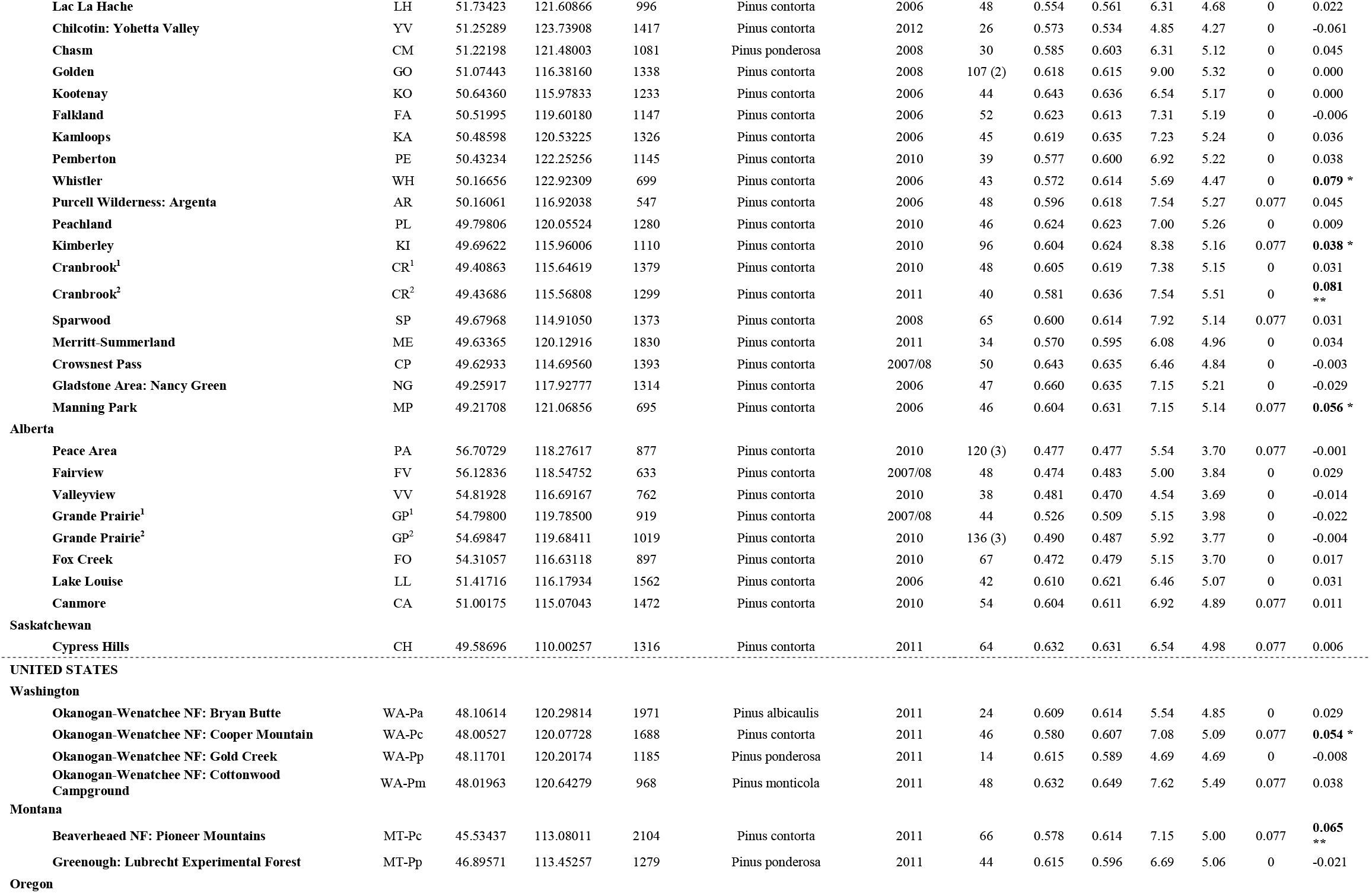
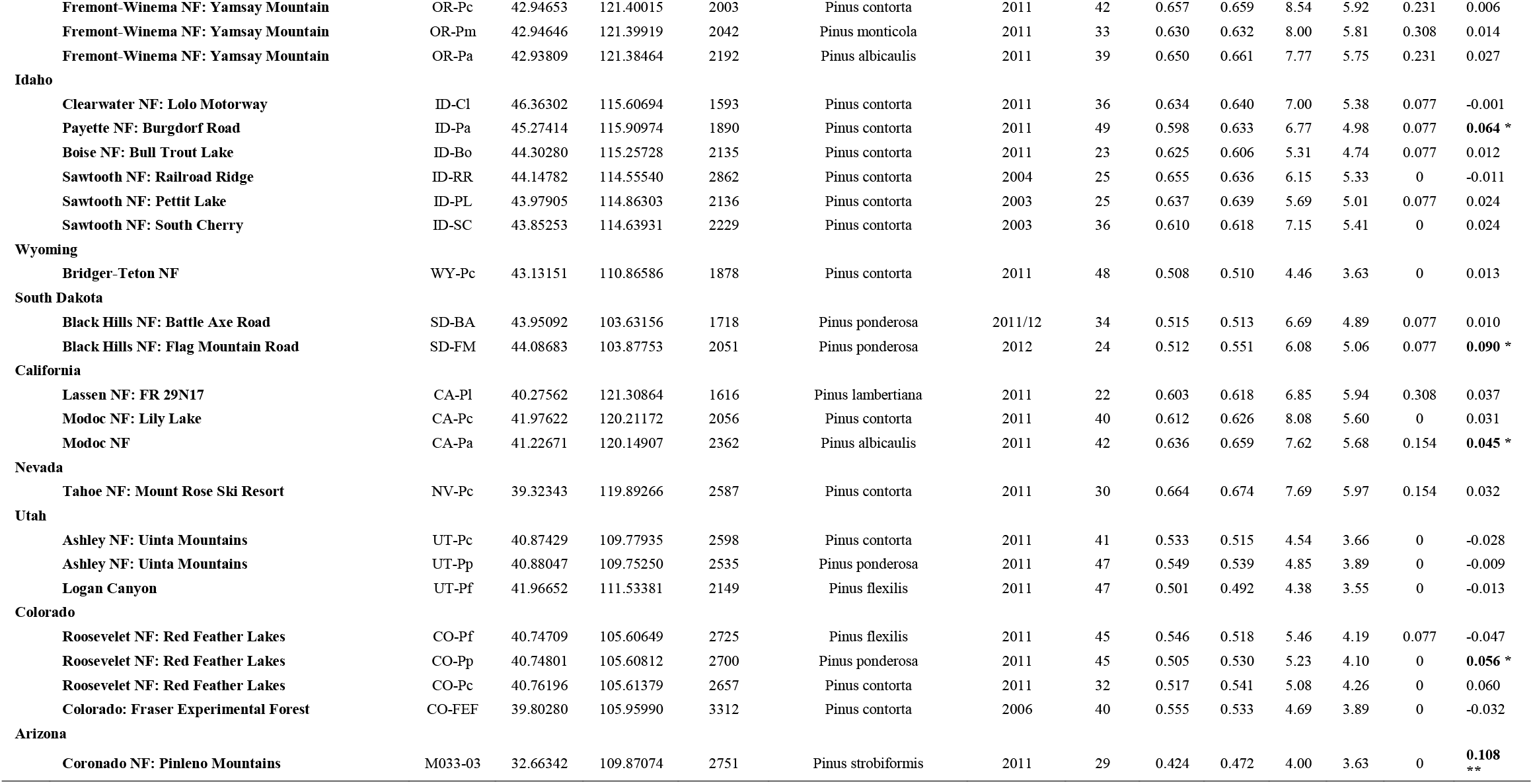
Sampling locations (82) by country and province/state for the mountain pine beetle. Variables shown include GPS coordinates, elevation, beetle host species, flight year sampled, number of beetles genotyped (N) (bracketed number indicates number of sites per location if multiple collections conducted). Mean observed heterozygosity (**H**_**o**_), mean expected heterozygosity (**H**_**e**_), mean number of alleles (**N**_**A**_), allelic richness (**A**_**R**_), mean number of private alleles (), and inbreeding coefficieint (**F**_**IS**_ - * p < 0.05, ** p < 0.005).

### DNA extraction and microsatellite amplification

Genomic DNA was isolated from whole beetles using a standard phenol/chloroform procedure (Sanbrook and Russell 2001) or an UltraClean Tissue and Cells DNA isolation kit (Mo Bio Laboratories, Carlsbad, CA, USA) following manufacturer’s protocol. Isolated DNA was resuspended in Tris-EDTA (pH 8.0) or eluted in the manufacturer’s buffer. Preliminary assessment of DNA quantity and quality was performed using a NanoDrop^®^ ND-1000 UV-Vis Spectrophotometer (Thermo Scientific, Waltham, MA, USA).

These analyses included a total of 3858 beetles genotyped at 13 microsatellite loci used for previous analysis within western Canada (Samarasekara et al. 2012), using four co-amplification procedures (Davis et al. 2009). Amplified fragments were co-loaded into two injections on an AB 3730 DNA analyser. Band sizes were determined relative to GeneScan-500 LIZ (AB) and scored using GeneMapper software.

### Hardy-Weinberg equilibrium and linkage disequilibrium

Genotypic data from each site were examined for Hardy-Weinberg Equilibrium (HWE) across loci and sites using an expansion of Fisher’s exact test. To ensure that all loci were independently assorting at all sites, linkage disequilibrium (LD; Slatkin and Excoffier 1996) was assessed using a likelihood ratio test. Statistical significance was evaluated both before and after sequential Bonferroni correction for multiple tests (Holm 1979; Rice 1989). All analyses were performed using ARLEQUIN 3.5.2.2 (Excoffier and Lischer 2010).

### Genetic diversity

Gene diversity and allelic richness were used to describe patterns of genetic diversity across the study area. Observed (H_O_) and expected (H_E_) heterozygosity were also calculated for each sample location using the PopGenKit package for R (Paquette 2012). Allelic richness was corrected for variation in sample size through rarefaction (Petit et al. 1998) executed in FSTAT 2.9.3.2 (Goudet 2001). Patterns of genetic diversity were assessed for the entire study area as well as within the main clusters identified by Bayesian analysis for population structure as described below.

### Population structure

Population genetic structure was determined using a Bayesian approach with STRUCTURE 2.3.4 (Pritchard et al. 2000), assuming an admixture model and correlated allele frequencies, without prior sampling information. Each run was performed with 100 000 burn-in and 50 000 MCMC steps with all other parameters maintained at default values. Population structure was tested at K ranging from 1 to 10 with 20 replicates. To assess membership proportions for clusters identified by STRUCTURE, the results of the 20 replicates at the best fit K were post-processed using Clumpp (Jacobson and Rosenberg 2007) and Structure Harvester (Earl and vonHoldt 2012). To capture a hierarchical level of population structure, each cluster was subsequently analysed for nested sub-structures and evaluated as previously described. Population structuring was visualized using ArcGIS version 10.5. To gain further insight into the underlying population genetic structure and provide additional support to the results generated using STRUCTURE, data were analysed within the framework of a discriminant analysis of principle components (DAPC; Jombart et al. 2008).

### Genetic differentiation

Genetic variance was partitioned among and within clusters using analysis of molecular variance (AMOVA) performed in ARLEQUIN 3.5.2.2 (Excoffier and Lischer 2010) based on pairwise F_ST_ corrected for unequal sample size using the method of Weir and Cockerham (1984). Variance components were calculated among groups (F_CT_), among locations within groups (F_SC_), among locations (F_ST_) and within sampling locations (F_IS_). Each AMOVA was run with 10 000 permutations at 0.05 significance levels.

To test portioning of variation associated with host use, non-lodgepole pine sites were paired with the closest (< 200 km) lodgepole pine site(s), thus ameliorating the effects of geography on the AMOVA groups. The sites were then placed into two groups: lodgepole and non-lodgepole. Non-lodgepole sites in AZ and SD were not used due the absence of sampling in lodgepole sites within 200 km (no sites within 400 km).

## Results

Based on our samples from 82 sites were, all 13 microsatellite loci typed displayed a high amount of polymorphism with between 10-36 alleles/loci reported (Table 1). Reflecting the large number of alleles, average H_o_ was also high ranging from 0.288 to 0.806. Similar to Samarasekara et al. (2012), no evidence for linkage disequilibrium among loci was found. Further, no sites displayed a large number of loci out of Hardy-Weinberg equilibrium (HWE), with each locus displaying a significant departure from HWE (p < 0.05) between 0-9 times out of the 82 sties tested. Significant positive F_IS_ values (p < 0.05) were noted for 16 populations with an overall F_IS_ value of 0.019 (p < 0.001).

Locus specific F_ST_ values were all significant and ranged from 0.040 to 0.219 (Table 2). This evidence of range-wide genetic structure is also noted in the overall F_ST_ value of 0.0683 (p < 0.001) among all 82 sampling locations. Pairwise population F_ST_ values range from nonsignificant to a maximum of 0.431 between sites in Arizona and northern Alberta (Figure 1). An outbreak in sugar pine (*Pinus lambertiana*) was also noted for its pairwise divergence (Fst 0.404 – 0168) from all other sites. Beetles from a successful brood noted in interior hybrid spruce (*Picea glauca X engelmannii*, McKee et al. 2015) were found to not significantly different from surround lodgepole pine sites in northern BC.

**Table 2.** Loci typed. Total number of alleles (N_A_), mean expected heterozygosity (H_e_), mean observed heterozygosity (H_o_), number of loci deviated from HWE, and fixation indices F_IS_, F_ST_ and significance values. *** = *P*< 0.0001.

| Locus | $N_A$ | $H_e$ | $H_o$ | Fixation indices | | | | |
| --- | --- | --- | --- | --- | --- | --- | --- | --- |
| | | | | HWE | $F_{IS}$ | $P$ | $F_{ST}$ | $P$ |
| Dpo028 | 25 | 0.488 | 0.463 | 7 | 0.051 | ns | 0.067 | *** |
| Dpo103 | 25 | 0.806 | 0.809 | 5 | -0.004 | ns | 0.053 | *** |
| Dpo160 | 36 | 0.717 | 0.723 | 7 | -0.008 | ns | 0.077 | *** |
| Dpo453 | 26 | 0.659 | 0.624 | 7 | 0.053 | ns | 0.039 | *** |
| Dpo479 | 11 | 0.612 | 0.607 | 4 | 0.008 | ns | 0.110 | *** |
| Dpo530 | 10 | 0.643 | 0.638 | 4 | 0.008 | ns | 0.061 | *** |
| Dpo566 | 12 | 0.288 | 0.293 | 3 | -0.016 | ns | 0.040 | *** |
| Dpo760 | 22 | 0.591 | 0.591 | 1 | 0.000 | ns | 0.076 | *** |
| Dpo780 | 15 | 0.564 | 0.563 | 0 | 0.002 | ns | 0.064 | *** |
| Dpo793 | 12 | 0.567 | 0.570 | 2 | -0.006 | ns | 0.128 | *** |
| MPB011 | 15 | 0.545 | 0.525 | 1 | 0.037 | ns | 0.048 | *** |
| MPB017 | 23 | 0.443 | 0.439 | 3 | 0.009 | ns | 0.168 | *** |
| MPB038 | 18 | 0.400 | 0.403 | 9 | -0.008 | ns | 0.219 | *** |

**Figure 1:**
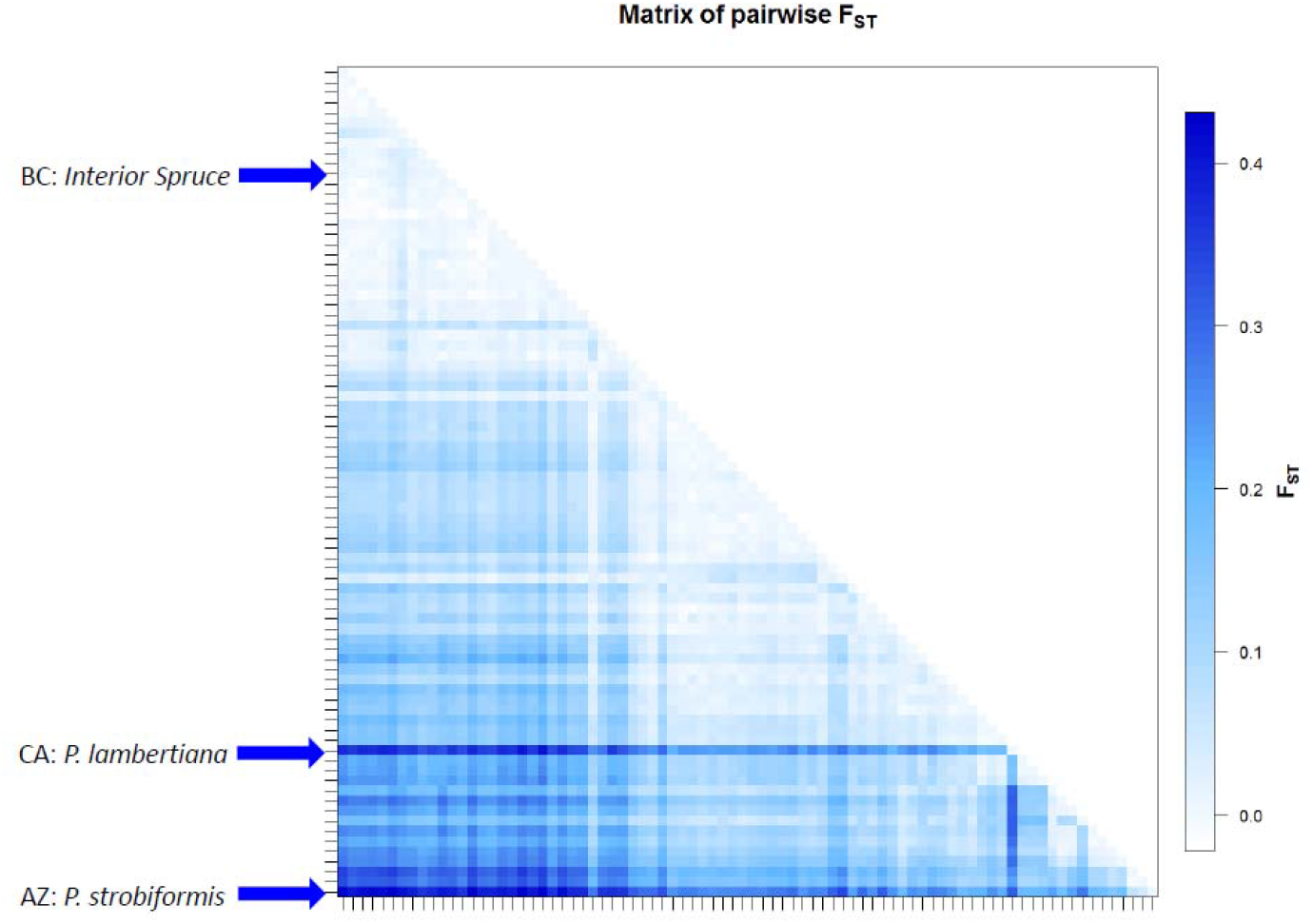
Pairwise Fst Matrix of all sites sampled.

To test the effects of host use on the observed genetic structure, subsets of the data were divided into 2 groups: lodgepole and non-lodgepole hosts (Table 3), covering a similar geographic range (Figure 2). In the first test, 14 non-lodgepole sites were paired with 11 lodgepole sites. The overall F_ST_ and F_SC_ were similar to each other and showed values consistent with the geographic range of sites examined. In contrast, F_CT_ values, the percent variation due to host use, is not significantly different from zero. Grouping the host site to test only: ponderosa pine and their closest lodgepole sites (5 pairs), white bark pine (4 pairs), and all others (2 western white pine, 2 limber pine and 1 sugar pine), all showed similar results.

**Table 3.**
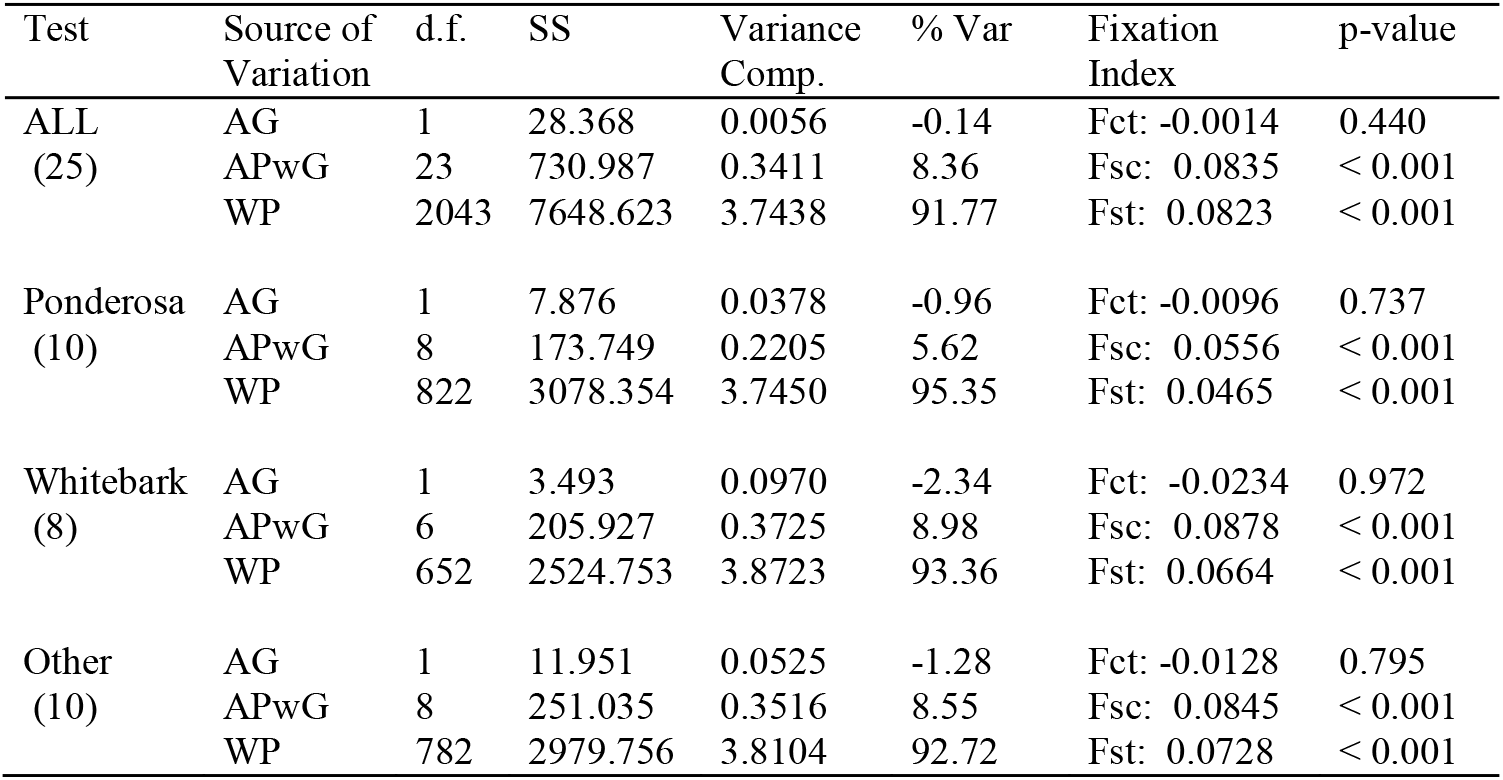
AMOVA analysis of population structure due to host use. Four tests performed including ALL combined non lodgepole, Ponderosa pine, Whitebark pine, and other pine test stands. Source of Variation includes AG (among groups), APwG (among populations within groups), WP (within populations). Shown for each source of variation are degrees of freedom, (d.f.), sums of squares (SS), variance component (Variance Comp.), percent of variation (% Var) and associated fixation index and p-value.

| Test | Source of Variation | d.f. | SS | Variance Comp. | % Var | Fixation Index | p-value |
| --- | --- | --- | --- | --- | --- | --- | --- |
| ALL<br>(25) | AG | 1 | 28.368 | 0.0056 | -0.14 | Fct: -0.0014 | 0.440 |
|  | APwG | 23 | 730.987 | 0.3411 | 8.36 | Fsc: 0.0835 | < 0.001 |
|  | WP | 2043 | 7648.623 | 3.7438 | 91.77 | Fst: 0.0823 | < 0.001 |
| Ponderosa<br>(10) | AG | 1 | 7.876 | 0.0378 | -0.96 | Fct: -0.0096 | 0.737 |
|  | APwG | 8 | 173.749 | 0.2205 | 5.62 | Fsc: 0.0556 | < 0.001 |
|  | WP | 822 | 3078.354 | 3.7450 | 95.35 | Fst: 0.0465 | < 0.001 |
| Whitebark<br>(8) | AG | 1 | 3.493 | 0.0970 | -2.34 | Fct: -0.0234 | 0.972 |
|  | APwG | 6 | 205.927 | 0.3725 | 8.98 | Fsc: 0.0878 | < 0.001 |
|  | WP | 652 | 2524.753 | 3.8723 | 93.36 | Fst: 0.0664 | < 0.001 |
| Other<br>(10) | AG | 1 | 11.951 | 0.0525 | -1.28 | Fct: -0.0128 | 0.795 |
|  | APwG | 8 | 251.035 | 0.3516 | 8.55 | Fsc: 0.0845 | < 0.001 |
|  | WP | 782 | 2979.756 | 3.8104 | 92.72 | Fst: 0.0728 | < 0.001 |

**Figure 2:**
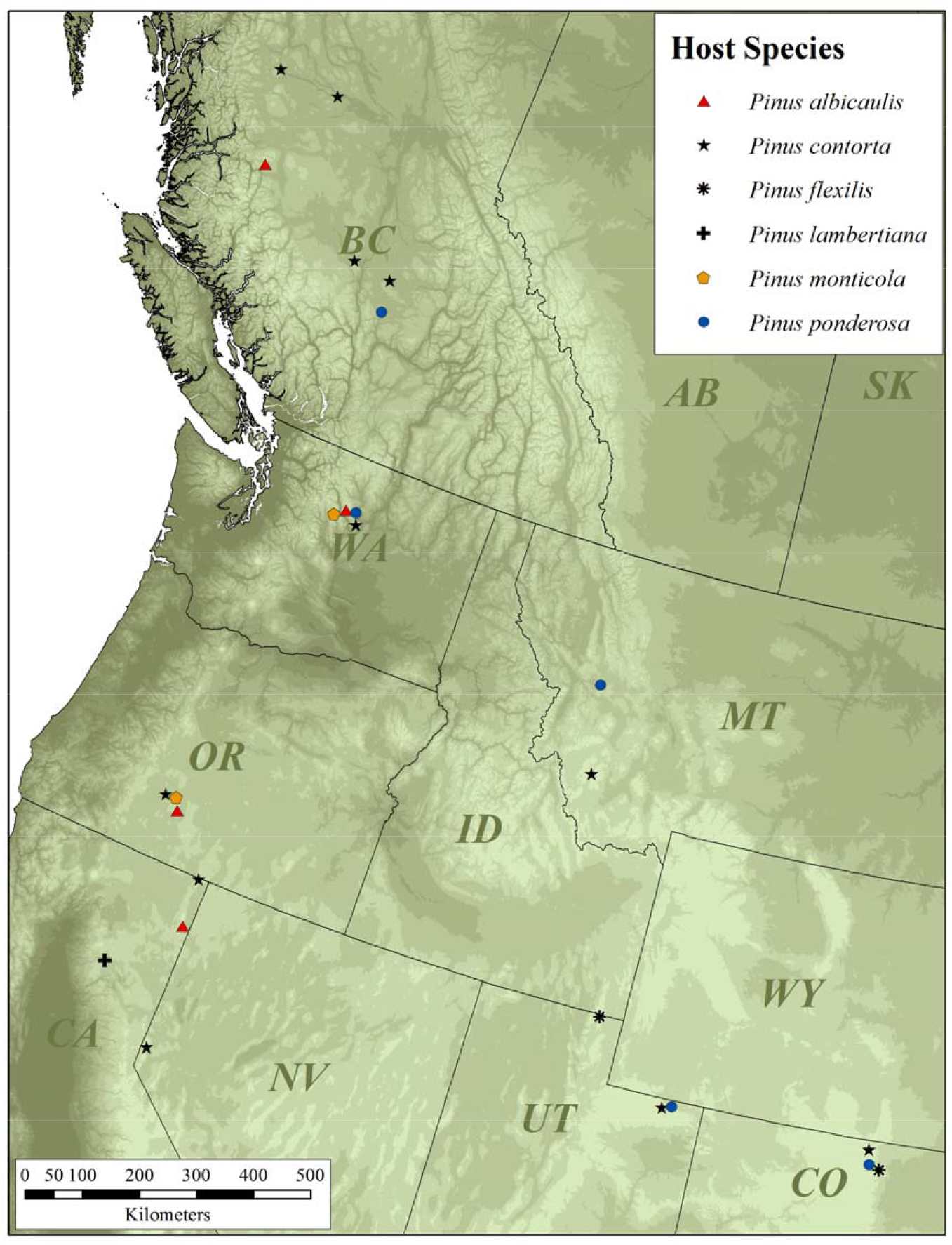
Map of study area with collection sites separated by host tree species. Closely located site markers were displaced from their geographic location on the map to allow for better visibility.

All 82 sites were used to examine range wide population structure. STRUCTURE analysis found two well supported clusters at K=2 dividing the samples into northern and southern groups (Figure 3). Among southern cluster, additional sub-clustering was found at k=3 and 4, supporting four clusters; 1 Northern - Northern BC/AB; 2 Central - Southern BC/AB/WA/ID/MT; 3 Southwest - OR/CA/NV and; 4 Southeast - UT/WY/AZ/CO/SD. This geographic population genetic structure was also found with DAPC analysis (data not shown).

**Figure 3:**
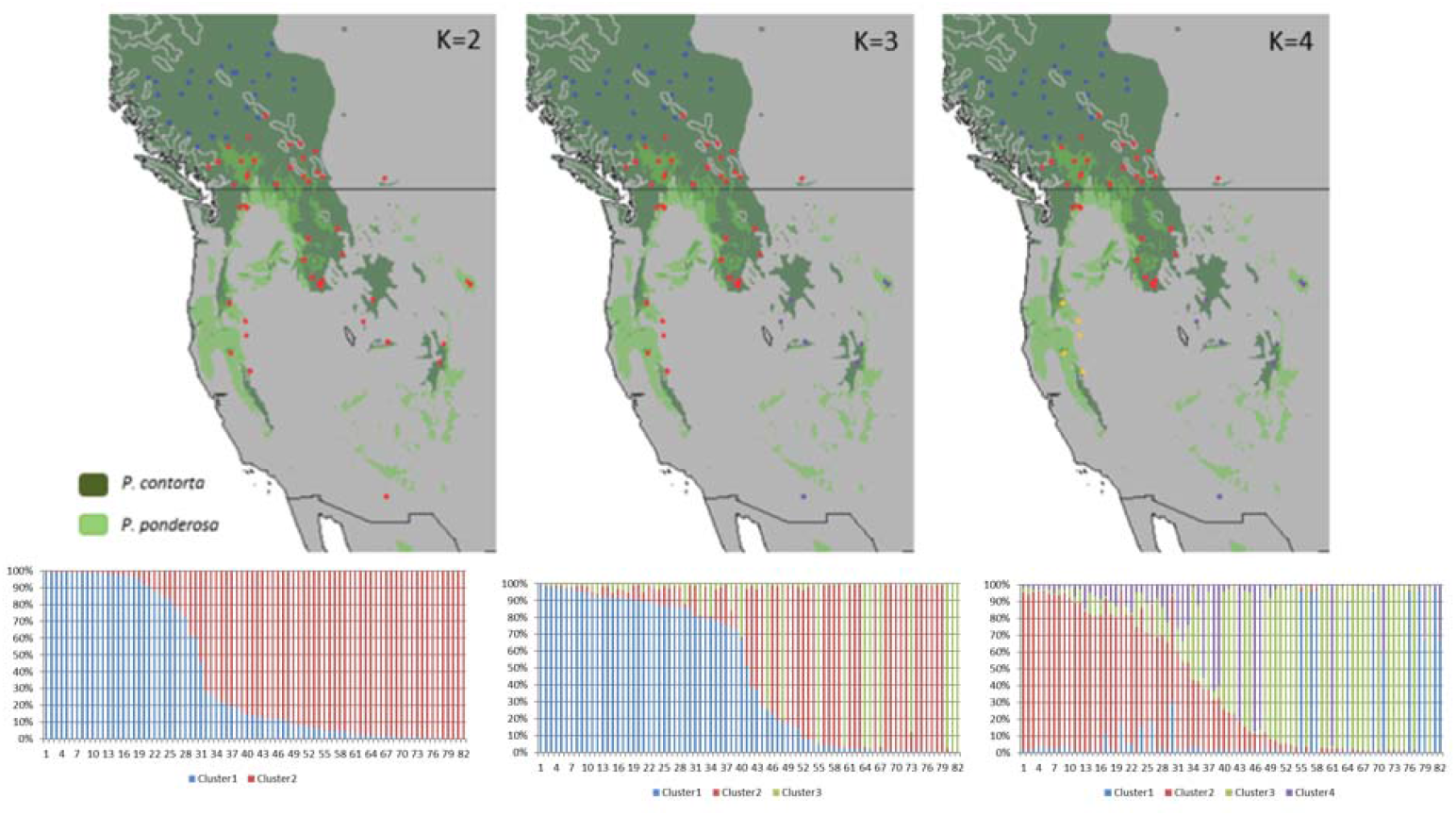
STRUCTURE analysis of 82 sites in western North America. K=2 is most strongly supported with weaker sub clustering, K=4, in the southern sampling locations. Distribution ranges for *Pinus contorta* and *P. ponderosa* are also shown.

Patterns of genetic diversity varied among the 4 genetic clusters. Of the 82 sites typed, higher allelic richness and heterozygosity values were detected in sites grouped into the southwest cluster (Figure 4). Both heterozygosity and allelic richness decreased in locations further north and to the southeast. The maximum H_O_ detected was 0.664 (Tahoe NF: Mount Rose Ski Resort) and the minimum was 0.424 (Terrace). The maximum allelic richness detected was 5.97 (Tahoe NV: Mount Rose Ski Resort) and the minimum was 3.45 (Terrace). The sites within the southwest cluster also showed the greatest number of private alleles (Table 1). The least amount of genetic variability was found in northern cluster populations compared to the rest of the data set (Figure 4).

**Figure 4:**
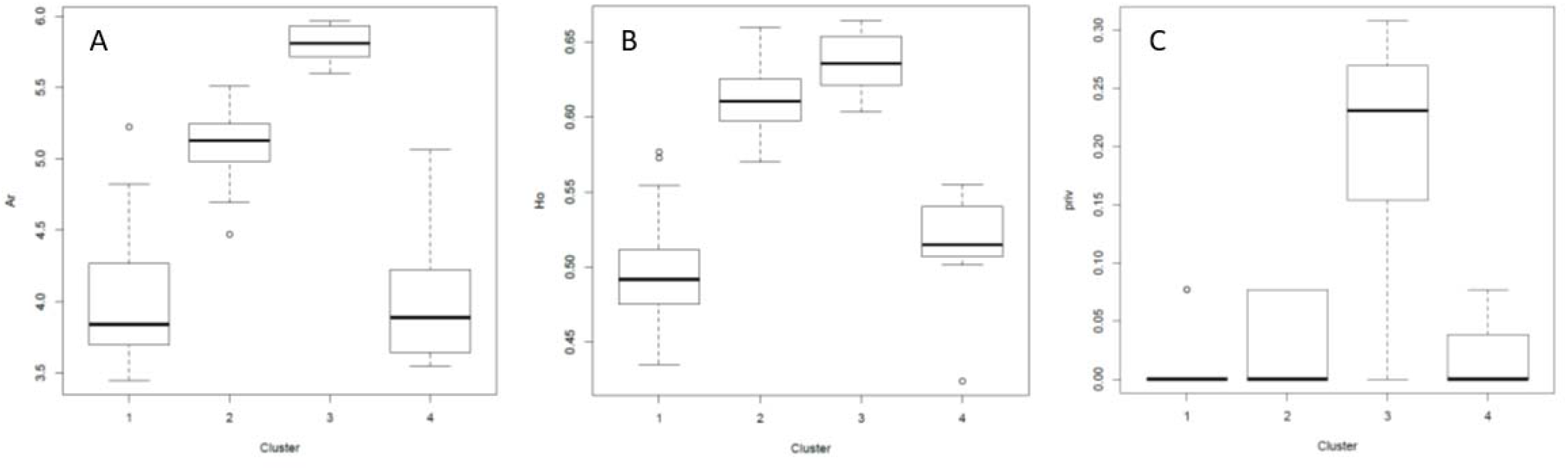
A) Allelic richness (Ar), B) and observed Heterozygosity (H_o_), and C) Private alleles grouped by STRUCTURE-generated cluster: 1 Northern - Northern BC/Alberta; 2 Central - Southern BC/Alberta/Washington/Idaho/Montana; 3 Southwest - Oregon/California/Nevada and; 4 Southeast - Utah/Wyoming/Arizona/Colorado/South Dakota. The boxplots incorporate Tukey’s 5-number summary as follows: the sample minimum, defined by the horizontal line at the bottom of each plot; the lower quartile, defined by the lower limit of the box in each figure; the sample median, represented by the heavy line inside each box; the upper quartile, defining the upper limit of each box; and the sample maximum, defining the horizontal line at the top of each plot. Open circles represent potential outliers falling outside the interquartile range.

## Discussion

### Host Use

Bark beetles exhibit behavioural plasticity in orientation sequences to various host species, which allows them to adjust to environmental variability, and are likely modulated by genetic, environmental, and genetic-X-environmental drivers (Raffa et al. 2016). For example, differential response to host phytochemical cues of third generation beetles of *Ips pini* (Scolytinae: Curculionidae) derived from breeding lines from endemic and irruptive populations has been shown to have a heritable component, specifically in host acceptance and gallery development (Wallin et al. 2002). Furthermore, only *Dendroctonus rufinennis* (Scolytinae: Curculionidae) beetles from eruptive populations tend to colonize well-defended, healthy spruce trees. While endemic and irruptive beetles may vary in their response to secondary defensive compounds, chemical similarities between historical and novel or non-traditional hosts can facilitate host range expansion (Erbilgin et al. 2013), and, hence, geographic range expansion of MPB. Early studies of natural infestation of various hosts species, e.g. Furniss and Schenk (1969), speculated that host utilization, including that of non-related species, i.e. *Pinus* and *Picea*, by irruptive MPB beetles may have aided in speciation of *Dendroctonus*.

Early allozyme studies showed genetic and morphological differences among preemergent adult beetles in different host tree species within the same stands suggesting potential incipient host-race formation (Sturgeon and Mitton 1986). In this analysis of five mixed hosts stands, they found slight allele frequency differences among hosts at 2 of 5 loci for all sites combined and inbreeding-like effects within stands, which they interpet as possible evidence of Wahlund’s effect between host species at a single site. Langor and Spence (1991) also report inbreeding-like effects at allozyme loci within sites and genetic differences among pre-emergent beetles among host species at a single site, but argue that the observed differences are due to differential survival/selective environments among hosts. Similar, less pronounced, results were observed among trees of the same host species within stands. Further, when the selective environment was reduced by collecting “slabs” and allowing beetle emegence, no genetic difference among host species was detected. These studies show that host tree condition/species can act as a strong selective force generating heterogentenity within MPB. Its potential role in the generation of host-races formation however depends on the fidelity of offspring to host species (Langor and Spence 1991).

Inbreeding-like effects were also noted in our analysis of neutral microsatellite markers. An overall F_IS_ of 0.019 and approximately 20% of sites showed significant F_IS_ values (Table 1). Significant F_IS_ values may be partially explained by the propensity of MPB to engage in sib-mating in the gallery prior to emergence (Bleiker et al. 2013; Janes et al. 2016). The enhanced inbreeding-like effects noted in allozyme studies may also be explained by the intensive sampling regime used in these earlier studies of primarily endemic populations. Although care was taken to sample as many galleries as possible, it is likely a large number of sibs would have been sampled given that Langor and Spence (1991) report that all available beetles in an infested tree were sampled. Genetic differences among hosts within a site may therefore reflect the differential reproductive success of individual broods within a single population (Langor and Spence 1991).

In contrast to the earlier studies of host use, our study analyses a single beetle per gallery and examines differences over the geographic range of the species. If host-race development is occurring, a signature of genetic difference among host should be detected at this larger scale. We find no evidence of genetic difference by host species and demonstrate the importance of geography to explain the genetic structure of MPB, as indicated in previous studies using several different marker systems (e.g. Mock et al. 2007, Samarasekera et al. 2012, Batista et al. 2016). The emergence of host-races requires host fidelity. A review by Barron (2001) suggests growing evidence of Hopkins’ host-selection principle, initially proposed in 1916, which refers to the observation that many adult insects demonstrate a preference for the host species on which they themselves developed as larvae. A study comparing MPB mating and host acceptance behaviour in a traditional host, lodgepole pine, and non-traditional host, Englemann spruce, showed that females reared in spruce more readily accepted spruce host material relative to pine, and had higher rates of host acceptance of both pine and spruce host material than females that had developed in pine (McKee et al. 2015). Monoterpene content of food sources of female MPB has been shown to influence the feeding choice of offspring (Burke and Carroll 2017). Although monoterpene content does vary with host, large variation within individual tree defensive capability is also found (Powers 1995; Huber et al. 2004). In contrast, the opportunistic nature of MPB host use is shown by their utilization of spruce during peak epidemic levels (McKee et al. 2015), in which we have shown MPB to be genetically identical to the surrounding lodgepole infections, and their expansion into boreal jack pine forests (Cullingham et al. 2011) during the recent outbreak in western Canada. Thus, although evidence exists for Hopkins’ host-selection principle in MPB, this may be balanced by host switching due to depleting ephemeral resources or host accessibility (i.e., wounded trees) leading to no detectable effect of host use on the range-wide genetic structure.

A possible exception to this trend was observed at a single sugar pine site examined in CA. This site was noted as an outlier in pairwise F_ST_ comparisons (Figure 1). While the range of pairwise values fall within that observed among other sites, it becomes notable when compared with other sites in CA. Curiously, the sugar pine has similar geographic range to Jeffery pine, which is infested by the host specialist Jeffery pine beetle. Because only a single site was examined, additional sites would be required to investigate the possible effects of host use in this species.

### Landscape Level Genetic Structure

Our expanded dataset presents strong evidence for clustering of MPB populations into two main groups, southern and northern, consistent with previous studies (Samarasekera et al. 2012; Cullingham et al. 2012; Janes et al. 2014; Batista et al. 2016). The northern cluster is particularly genetic homogenous, showing comparative decreases in heterozygosity and allelic richness. This is consistent with the expectations of founder effect during range expansion (Allendorf et al. 2012) and where mating practices are more random, and traits are not exposed to the same level of environmental selection due to the number of new mutations and lineages within the expanded population (Klopfstein 2005). While MPB is not truly panmictic, even as an airborne insect, it is likely that environmental homogeneity of northern BC has facilitated more interbreeding than the southern populations that are separated by heterogenous habitat and landscape barriers, such as mountains.

Our work also shows sub-clustering in the southern cluster which is similar to that reported from mitochondrial (Cullingham et al. 2012) and SNP datasets (Batista et al. 2016; Dowle et al. 2017). Differences among the studies may be partially explained by the resolution of the different marker systems used and the distribution of sampling locations. Batista et al. (2016) have a high number of sampling locations in Canada with relatively few in US (n=10), while Dowle et al. (2017) has the majority of sampling locations in the US and few in Canada (n=2). Our study attempted to even out the geographic distance among locations, but still lacks important regions in the US sampled by Dowle et al. (2017). A general picture emerges of a central cluster extending from southern BC into the US states of ID and MT. Locations from Cascade mountain in WA cluster more strongly with this central cluster in our analysis, while SNP based studies place MPB collected in similar locations into a southwest cluster which extends southward into OR and CA. Evidence for additional southeast clusters also exists. Our study shows a southeast cluster breaking from the central cluster at the snake river plain including locations from UT/WY/CO/AZ/SD. Analysis of autosomal SNPs in this region, which includes a greater number of sampling locations in AZ and central NV, reveals the presence of an additional cluster(s) - locations from AZ, NV and SD separate from those found in ID and CO (Dowle et al. 2017). Despite genetic evidence of gene flow among the clusters, mating studies have shown reproductive barriers among locations from the southwestern and central/south-eastern clusters which indicate possible incipient speciation (Bracewell et al. 2011; Bracewell et al. 2017).

### Indications of Glacial Refugia

Observed heterozygosity (H_o_) and allelic richness are both measures that are used as a proxy for expressing the genetic resilience and durability of populations (Bromham 2008; Caballero and García-Dorado 2013). When our study sites are partitioned by the 4 clusters generated by STRUCTURE, the southwest cluster from OR/CA/NV has the highest values for both H_o_, allelic richness and private alleles. High allelic richness is especially important as it provides a source for adaptive variation as populations face new pressures (Greenbaum et al. 2014). Previous research also found high genetic variability within the same geographic region, with low diversity in the northern range (Mock et al. 2007; Cullingham et al. 2012). This clinal reduction of variation in more northerly populations compared to southerly populations mirrors the glacial retreat from the last glacial period of the Quaternary (Konnert and Bergmann 1995; Schaetzl and Anderson 2005). It is likely that MPB populations followed the spread of pine northward during glacial retreat, but due to the restraint of the outbreak cycle, expanded more slowly than wind and animal assisted pine species (Burke et al. 2017).

The Pacific Northwest of the United States has previously been identified as a glacial refugia for many different species (e.g. Shafer et al. 2010). The high levels of diversity and unique alleles observed in the southwest cluster indicates the importance of this region as a reservoir of MPB genetic diversity. This region possesses a complex orogeny and is geographically heterogenous with many possible barriers to animal movement. It is important to note that, while MPB is capable of flight, it requires external forces such as wind to assist its movement over large land barriers, such as mountain ranges (Jackson et al. 2008; Evenden et al. 2014). This partitioning of habitat has led to the concept of refugia-within-refugia for the Pacific Northwest (Shafer et al. 2010). Although it is tempting to view the observed pattern of reduced diversity as a pathway of post glacial expansion (i.e., from the southwest glacial refugia to the central cluster, then north and southeast), an analysis of neo-Y haplotype diversity shows a more complicated pattern of post glacial expansion. Dowle et al. (2017) find three geographically separated neo-Y haplotype lineages; southwest (CA/OR/WA), southeast (AZ/NV/SD) and the Rocky Mountains (CO/UT/WY/ID/MT/BC/AB) indicating 3 core refugia. It is likely that the genetic diversity in the central cluster increased relative to the founder population due to gene flow of autosomal markers from the other main refugia while at the same time maintaining Rocky Mountain neo-Y markers due to reduced hybrid male fertility (Bracewell et al. 2017). In this light, northern post glacial expansion into Canada was primarily from a refugia associated with the rocky mountain neo-Y lineage, but likely also carried an autosomal legacy from all three refugia.

## Conclusions and Future Directions

Results from the dataset used in this study suggest that host tree species of MPB has not contributed to the genetic structure of populations to a detectible level, an important finding which refutes the hypothesis of the development of cryptic host-races of MPB. The lack of influence of host species indicates that forestry personnel are better served making management judgments based not on species of attacked trees, but instead on stand structure and composition, ecological and environmental site conditions, and beetle population phase and brood success.

This study also illustrates the complexity of interpreting spatial genetic patterns in terms of post-glacial expansion. Although the Pacific Northwest refugia likely represented a significant reservoir of genetic diversity, it is unclear how important this refugium was in the northern expansion into Canada. The wider genomic distribution of SNP datasets, including neo-Y coverage, make these markers more suitable to address this question. Future studies should examine populations across the contact zone of the neo-Y haplotypes as identified by Dowle et al. (2017) within WA and southern BC for both gene flow and mating success (i.e., Bracewell et al. 2017) to more fully understand the genetic contribution of the refugia populations to the MPB of WA, ID, MT and southern BC. Further, these studies can then be used to inform studies on the genetic makeup of the historic and current range expansion of the northern population.

## Acknowledgements

The researchers wish to thank the many people who aided with sample collection (see Table S1). Thanks to Dr. Neil Thompson for his detailed GIS assistance. Special thanks to Sophie Dang for her excellent microsatellite typing work. We are grateful for the field and laboratory assistance of Rose Loerke, Shane Doddridge, Mike Prior, Rhiannon Montgomery, Cierra Hoerchl, Will Eisbrenner, Amanda Cookhouse, and Marcelo Mora. Funding for various aspects of this project was provided by a Genome Canada, Genome BC, Genome Alberta and an NSERC Strategic Network Grant to the TRIA Project and by BC FLNRO, AB ESRD, AEP, SK Environment, USDA Forest Service, Utah State University, NV Department of Forestry, White Rose Ski Resort, NV.

## Notes

### Competing Interest Statement

The authors have declared no competing interest.

