## Supplementary figures and images for "Host use does not drive genetic structure of mountain pine beetles in western North America"

### Supplemental Figure 1

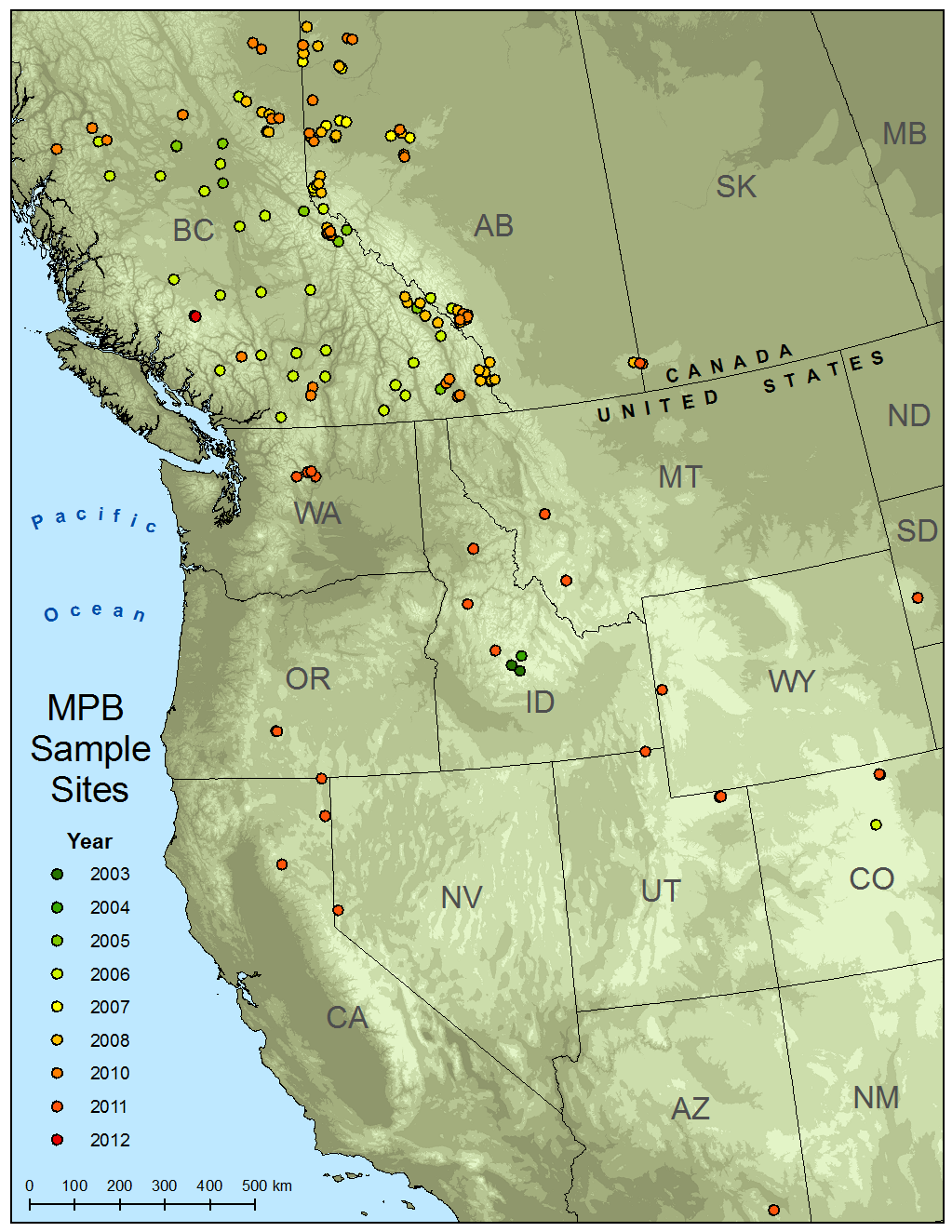

Supplemental Figure 1: Map of all 82 sites sampled for MPB in North America colour coded by year
