## Supplemental Table 1 for "Host use does not drive genetic structure of mountain pine beetles in western North America"

Supplemental Table 1: Collaborators for mountain pine beetle sample collections in western North America, 2011-2012.

| **CANADA** |  |
| --- | --- |
| **British Columbia** |  |
| Smithers | Ken White, Erin Havard (BC MFLFRO) |
| Terrace | Aaron Benterud (BC MFLFRO) |
| Mackenzie | Andrew Tait (BC MFLFRO) |
| Tumbler Ridge | Stephanie Haight (BC MFLFRO) |
| Fort St. John | Stephanie Haight (BC MFLFRO), Seth Folkman (Contractor) |
| Valemount | Steve Gillette (BC MFLFRO) |
| Chilcotin | Leo Rankin (BC MFLFRO), Joe Cortese (Contractor) |
| Kimberley/Cranbrook | Liz Goyette (BC MFLFRO), George Eimer (Contractor) |
| Peachland | Wolfgang Beck (BC MFLFRO) |
| Merritt | Cliff Robertson (Tolko Industries Ltd.) |
| Whistler | Andrea Lyall, Tom Cole, Stirling Angus (BC MFLFRO) |
| **Alberta** |  |
| Kananaskis/Spray Lakes | Brad Jones (AB ESRD), Rod Gow (AEP) |
| Fox Creek/Valleyview | Anina Hundsdoerfer (AB ESRD) |
| Grande Prairie | Devin Letourneau, Pam Melnick (AB ESRD) |
| Peace Area | Lindsay Eastman (AB ESRD) |
| **Saskatchewan** |  |
| Cypress Hills | Rory McIntosh, Robert Moore, Jeff Gooliaff (SK Environment) |
| **UNITED STATES** |  |
| Region 1: Northern | Brytten Steed (MT); Lee Pederson (Northern ID) |
| Region 2: Rocky Mountain | Bill Schaupp, Jim Blodgett (SD); Sheryl Costello (CO) |
| Region 3: Southwestern | Andrew Graves (NM); Joel McMillan, Ann Lynch (AZ) |
| Region 4: Intermountain | Carl Jorgensen (southern ID); Steve Munson, Ben Meyerson (UT, western WY); Barbara Bentz, Matt Hansen (UT);  * Karen Mock (Utah State University); * Gail Durham (NV Department of Forestry)  * White Rose Ski Resort, NV |
| Region 5: Pacific Southwest | Daniel Cluck (northern CA) |
| Region 6: Pacific Northwest | Andrew Eglitis (OR); Darci Carlson (WA) |
